# Integrating Safety, Security, Sustainability and Social Responsibility Principles into the U.S Bioeconomy

**DOI:** 10.1101/2024.02.05.578954

**Authors:** Aurelia Attal-Juncqua, John Getz, Ryan Morhard, Gigi Kwik Gronvall

## Abstract

Bioindustrial manufacturing is undergoing rapid expansion and investment and is seen as integral to nations’ economic progress. Ensuring that bioindustrial manufacturing benefits society as the field expands is of critical, urgent importance. To better understand the industry’s ethical trajectory and to shape policy, we explored the views of biotechnology leaders on 4 aspects of ethical and social responsibility: - Safety, Security, Social Responsibility, and Sustainability, what we have termed “4S Principles”. We identified policy actions governments and other stakeholders may take to maximize societal benefits in industrial biotechnology.

## Main Text

Biomanufacturing, a sector devoted to producing goods derived from biological processes (from pharmaceuticals and biofuels to bioplastics and other biomaterials), serves as a cornerstone of the broader bioeconomy, creating a tangible link between innovation and market-driven production of biological products. The US bioeconomy—currently valued at more than $950 billion and accounting for more than 5% of US gross domestic manufacturing—is growing rapidly with increasing impact on the country’s economic vitality.(*1*) As biomanufacturing continues to expand, its profound and wide-ranging influence on society heightens the importance of integrating ethical considerations into the industry. While initiatives like the National Institutes of Health (NIH)’s Ethical, Legal, and Social Implications (ELSI) Research Program have addressed biotechnology’s societal and medical impacts, growing global competition and climate change highlight the need for broader societal considerations, including sustainability and social responsibility.(*2*) Bioindustrial manufacturing offers an opportunity to incorporate these societal norms as the field continues to progress on the global stage, with new models for how they can be incorporated into technology development.

As an example of this, the Bioindustrial Manufacturing and Design Ecosystem (BioMADE), established by the US Department of Defense (DoD) in 2020, has prioritized incorporating “4S principles”—Safety, Security, Sustainability, and Social Responsibility—into the biomanufacturing sector (table 1), and into all technical projects funded by BioMADE.(*3*)

**Table 1.** 4S Definitions.

| Principle | Definitions |
| --- | --- |
| Safety | Practices, controls, and measures taken to protect people and the environment from harm from biomanufacturing development processes and/or physical product or byproducts. Includes safety of the workplace, consumers, and the general public. |
| Security | Measures taken across the biotechnology and biomanufacturing sectors, including food and agriculture, materials, and energy, to manage potential threats and loss due to theft, misuse, diversion, unauthorized possession of property (including intellectual property [IP]) or intentional release of biological risk and/or technology. |
| Sustainability | Measures taken to maintain or improve the long- |
|  | term viability of the environment and economy due to advancing biomanufacturing processes. These would include consideration of the impacts of products and processes on the environment, supply chain, as well as local public/consumer acceptance and practices. |
| Social Responsibility | A principle that acknowledges the impacts of biomanufacturing on stakeholders with respect to associated benefits, risks, and consequences throughout the value chain. This implies taking actions that optimize positive social outcomes through adherence to ethical standards, including seeking ways to make products and processes that improve societal welfare. Special attention to this commitment includes equitable distribution of benefits and risks and a responsiveness to society's needs and values. |

[source: Aurelia Attal-Juncqua, Galen Dods, Nicole Crain, James Diggans, David Dodds, Steve Evans, Nick Fackler, Kevin Flyangolts, Kathleen Gibson, Margaret E. Kosal, Aditya Kunjapur, Russ Read, Brian Renda, Corinne D. Scown, Kissaou Tchedre, Krista Ternus, Beth Vitalis, Gigi Gronvall, Shaping the future US bioeconomy through safety, security, sustainability, and social responsibility, Trends in Biotechnology, 2023]

This paper explores biotechnologists’ perceptions of the 4S principles and seeks to establish a foundational understanding of their views. By examining the opinions of industry experts, many of whom are affiliated with BioMADE, we aim to identify actions that can be taken to enhance the societal benefits emerging from bioindustrial manufacturing and to continue shaping future biomanufacturing policy more broadly. While interviewees were largely focused on actions the US could take, promoting 4S principles in bioindustrial manufacturing should be a priority all governments that are investing in the growth of this field.

## Methods

The Johns Hopkins University Bloomberg School of Public Health Institutional Review Board determined that this study did not constitute human-subjects research [IRB00023291].

### Interviews

From April 2023 to June 2023, the researchers conducted a series of semi-structured, virtual interviews with 31 industry leaders, representing a variety of perspectives from the BioMADE memberships. These leaders included individuals associated with academic institutions, as well as CEOs, founders, CSOs, and CTOs of biomanufacturing companies. The researchers developed an interview guide based on results of an informal conversation with BioMADE leaders, as well as the researchers’ personal experience and expertise related to biosafety and biosecurity. While the interview guide included core topics, interviewees were allowed to direct the conversation based on their individual experiences and priorities. All interviews were conducted on a not-for-attribution basis to promote candor and transparency. During each interview, a member of the research team took notes and audio was recorded and transcribed— with interviewees’ consent—to supplement interview notes.

### Analysis

The researchers employed a qualitative approach to analyze interview content, systematically and rigorously documenting the landscape of perceptions associated with BioMADE’s 4S Principles. Qualitative coding of interview notes was done using NVivo qualitative coding software and priority themes were identified and coded. The initial thematic coding framework was solely based on BioMADE’s 4S Principles. The researchers added themes as they emerged during the interviews. The final coding framework included: Sustainability, Safety, Security, Social Responsibility, Education, Public Risk Perception, Competition, Regulatory Space, and Workforce.

Using NVivo12 Pro and Microsoft Excel, semi-quantitative metrics were generated for all codes in the framework to measure the frequency with which they were discussed. These metrics included the number of coding references (individual chunks of coded text) corresponding to each code. Some references were co-coded. These descriptive metrics were also used to identify themes discussed more often or more in-depth, which could signal differences in how stakeholders prioritize certain topics. The research team also conducted a thorough qualitative analysis of the coded references, undertaking a detailed review of the coded text corresponding to highlighted codes, enabling the researchers to identify important comments and recommendations, both those that were prevalent across numerous interviews and those that were not.

### Findings

While the study was not designed to yield quantitative results, the findings reflect the relative frequency with which interviewees discussed topics, expressed viewpoints, or provided recommendations (e.g., “some” or “many” interviewees).

### Results Overview

Researchers invited 62 industry leaders to participate in this study, of which 31 were interviewed (table 2). Recruited participants came from a diverse range of academic and industry backgrounds. The majority are BioMADE members, but additional participants were identified through snowball sampling.

**Table 2.** Participant List.

| **Name** | **Title** | **Organization** |
| --- | --- | --- |
| Diggans, James | Head of Biosecurity | Twist Bioscience |
| Calder, Shasha | Head of Impact | Genomatica |
| Davis, Amy | Gov Relations | Novozymes |
| Kuldell, Natalie | Founder | Bio Builder |
| Herr, Daniel | Professor | UNCG |
| Matlock, Peter | Bioeconomy Research & Commercialization Specialist | University of Hawai'i at Hilo |
| Webber, Jo | CEO | STEMconnector |
| Wang Ben | Professor | Georgia Tech |
| White, Chip | Professor | Georgia Tech |
| Ajikumar, Parayi | Founder & CEO | Manus Bio |
| Carr, Peter | Senior Staff | MIT |
| Leproust, Emily | CSO / CEO | Twist Bioscience Corp. |
| Prather, Kristala | Chief Scientist | Kalion Inc |
| Bruno, Marilyn | CEO | Aequor |
| James, Joseph | President | Agri-Tech Producers, LLC |
| Franklin, Scott | Chief Scientific Officer | Checkerspot |
| Magyar, Andrew | Chief Technology Officer | Capra Biosciences |
| Demirel, Melik | Technical Advisor | Tandem Repeat Technologies |
| Hess, Mike | Sr. Manager - Global Optimization | Novozymes |
| Ternus, Krista | Senior Science & Technology Advisor | Signature Science |
| Tracy, Bryan | CEO | Superbrewed Food |
| Tyler, Christopher | Manager | Cargill |
| Garcia, Fernando | Senior Director, Scientific & Regulatory Affairs | Amarys |
| Tasseff, Ryan | Chief Technology Officer | Biocellion |
| Cameron, Doug | Advisor/Consultant |  |
| Carnstens, Kerri | Chief Executive Officer | Jordbioscience |
| Gray, Kevin | Biotechnology Executive | Kevin Gray Consulting |
| Barbero, Robbie | Chief Business Officer | Ceres Nanosciences |
| Sato, Aaron | Chief Scientific Officer | Twist Bioscience |
| Bitting, Angela | Chief ESG Officer & SVP, Corporate Affairs | Twist Bioscience |
| Starr, Jack | Director | Cargill |

The research team performed thematic coding on transcripts, resulting in 580 total coding references. The number of coded references for each of the 4S Principles were similar, with 22.4% of the references coded to Sustainability, 20% to Security, 16.6% to Social Responsibility, and 15.7% to Safety. The added codes (Education, Public Risk Perception, Competition, Regulatory Space, and Workforce) account, together, for 25.3% of the coded references (Fig. 1). Of note, the researchers found that 52.6% of the coded references to the “Security” Principle originate from only 4 interviews, while references coded to the 3 other Principles were evenly distributed across interviews.

**Figure 1.**
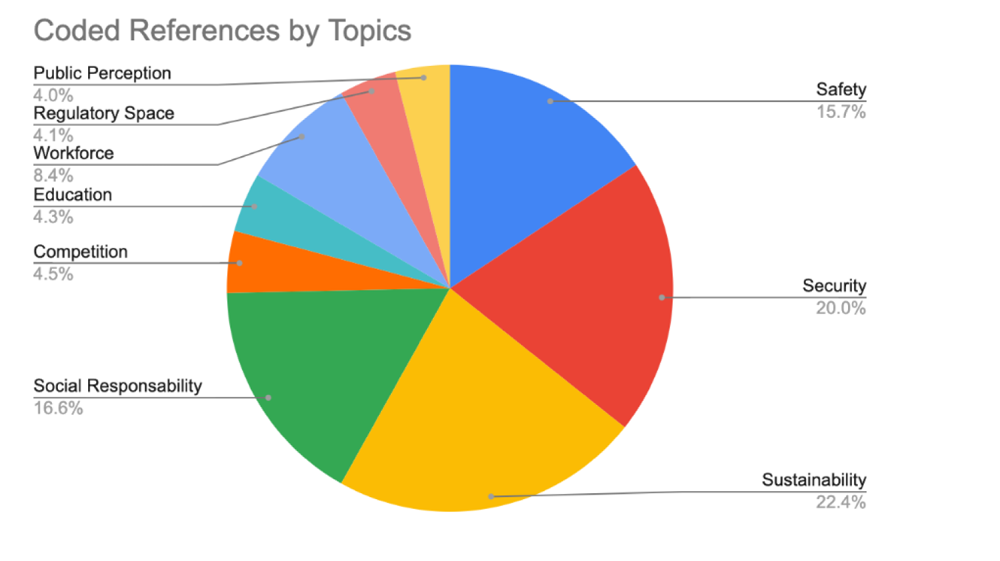
Descriptive Summary of Coded References.

### Participants’ 4S Perceptions

Participants discussed their understanding of each 4S concept. Most industry leaders exhibited a homogenous understanding of Safety, Sustainability and Social Responsibility, there was a diverse understanding of Security. Most stakeholders interpreted security concerns as revolving around IP, economic security, supply chain resilience, export control and cybersecurity. Only a minority expressed concerns about biosecurity, such as risks like AI-enabled creation of harmful products, toxins or pathogens (see table 3).

**Table 3.** Main Findings.

| **SAFETY** |  |  |  |
| --- | --- | --- | --- |
| **Perception** | **Barriers** | **Recommendations** | **Key Quotes** |
| Risk versus hazard evaluation<br>Biosafety procedures, best practices & training<br>Safety for human consumption, microbiomes & the environment<br>Personal & worker safety<br>Biocontainment & environmental controls (e.g., steel reactors)<br>Strong overlap with social responsibility | Risks assessments for de novo products is more complex to undertake<br>Patchwork regulation on engineered organisms (e.g., state by state approach for soil organisms)<br>Complicated regulatory landscape due to diversity in product type & usage<br>Public perception<br>Missing metrics and evaluation techniques | clear definition of the term, the goals and what success look like<br>Promoting safety culture<br>Studies to monitor adverse effects on environment & microbiome<br>Engage with the community<br>Consider a hierarchical approach to screening new materials as they progress from academia into prototyping into scale up manufacturing.<br>Explore “chemical reach across programs” for products that have “grandfather chemicals” which have already been tested for safety to even out the playing field and limit undue regulatory barriers<br>Develop and promote metrics and measurement techniques to determine whether a company is meeting standardized safety goals<br>Directly invest in research on the safety and security implications of new tech & build risk models<br>Increase government investment in developing /expert capacity<br>Need more national level cohesion (e.g., Biological Products Industry Alliance)<br>Support newer, smaller companies to navigate the fragmented regulatory landscape | How do we engineer out the risks?”<br>“The overarching sentiment in the industry is that there are already protocols in place to make sure that you have the right environmental control to minimize escape. But the risks are changing.”<br>“We lack the expertise to even determine when and how the risk may arise, and how to detect.”<br>“Safety regulations are completely inconsistent –so everybody gets stalled out”<br>“Bio-based ingredients must go through full toxicity analysis, sometimes animal testing and so forth. It's very costly, and it causes great delay”<br>“Bigger companies are better able to handle this patchwork approach because they actually have a regulatory team that can do that. That doesn't mean a little company can't. But it might be harder.”<br>“there's really terrible safety regulation uniformity on engineered organisms in the US.”<br>“I'm frustrated by the pace at which our safety and regulatory bodies are able to adapt to and keep pace with technological innovation”<br>“On the nuclear side, the national labs really took the lead on trying to explore safety and security risks. And I and I just don't see that kind of like large scale investment going into the National Lab architecture on the bio side.” |
| **SECURITY** |  |  |  |

| Perception | Barriers | Recommendations | Key Quotes |
| --- | --- | --- | --- |
| Intellectual Property concerns<br>Export control<br>Foreign staff<br>Public health, economic security, and defense (e.g., bioterrorism, including agricultural bioterrorism)<br>Gene synthesis & storage<br>Physical, industrial security (e.g., hacking events) - as the economy moves towards more biomanufactured products, these facilities have the potential to become targets<br>Cybersecurity<br>Domestic resilience/supply chains<br>Risks associated with AI enabled synthesis of de novo molecules | Lack of clarity on export control/ITAR requirement related to biotech<br>Lack of tools to screen for new functionality of de novo DNA, as existing screening tools rely on homology: the emergence of new AI tools that facilitate the creation of novel sequences not present in nature, or those with innovative functions, has rendered the screening process using current tools less effective, if not somewhat blind.<br>Lack of national level regulation for non-US-based synthesis company<br>Lack of regulation for US companies that buy DNA from outside of the US<br>No existing capacity or expertise to properly assess how different biological functions would interreact with each other in a single cellular system and identify potential risks<br>Lack of metrics or measurement techniques to assess whether organizations are successful at achieving set security goals | Share standardized clear definitions<br>Fund or undertake additional research to assess how different biological functions would interreact with each other crammed into a single cellular system<br>Support the development of metrics and measurement techniques<br>Gov and companies should work together on the development of clearer rules and guidance for companies<br>Develop foreign staff consideration guidelines or requirements<br>Explore export controls policies<br>Directly invest in research on the safety and security implications of new tech, build risk models.<br>Develop broadly available tools that estimate the risk and screen sequences<br>Require all US companies to purchase DNA from the verify provider<br>Hold/fund workshops with cross- sector representation to try to develop potential policy recommendations for government to address the risks surrounded AI derived proteins<br>Invest in the development and roll out of genetically built in IP protection with the ability to thwart an unauthorized user from being able to take advantage of an organism<br>Policy research and development related to economic security in the context of the transition to a bio-based economy (e.g., policy surrounding who buys corn futures in the United States), including research on the data infrastructure needed for geo-economic | "We need to better understand the implications of moving from a carbon-based economy more to a bio-based economy."<br>"In order to prevent them (competitors) to capitalize on our IP, we put genetic switches that they're not aware of, so that the strain will continue to refuse any new DNA. They're commercially being applied today and could be applied as bio terrorism protection mechanism."<br>"What do the systems do when put together? And is the outcome something that would be weaponizable?"<br>"I think right now there's kind of a chicken and egg problem. If the US government tomorrow said we're only going to spend our dollars with responsible companies. But who are the responsible companies? You can make an argument that it's the IGC companies, but like that's one level of a surety, but it's not a third-party level of a surety. It's really not anywhere near the equivalent of an ISO audit. What is the level of confidence that you would need to have before you're willing to redirect state, or Federal dollars to a limited set of economic actors?"<br>"Historically, the government had a responsibility to understand the safety and security implications of emerging technologies. I just don't see that kind of like large scale investment going into the National Lab architecture for bio."<br>"We just have no ability to model or predict negative outcomes on any biological level. It seems like we should be working on that, |
|  |  | forecasting.<br>Invest/ Support supply chain visibility – understand where your feedstock is coming from and what are the economic and security implication of that/ assess geopolitical risk of sourcing – learn from other industries how to better understand your own supply chain | since we're working furiously on the ability to design totally de novo biological functions."<br>"Because biotech is so fragmented, safety and security can mean different things to different people, and the significance of it can be different."<br>"If you're sponsored by the Department of Defense, you think about safety and security because you're forced to. If you are interested in more entrepreneurial work, you start to think about security from the perspective of intellectual property."<br>"There are things/risks that you don't know you don't know." |

SUSTAINABILITY
| Perception | Barriers | Recommendations | Key Quotes |
| --- | --- | --- | --- |
| Transitioning from fossil fuel-based economy to a bio-based economy<br>Climate change & Resilience<br>Economic Resilience<br>Environmental sustainability<br>Economic sustainability<br>Human health Impacts | Variance in data sets and methods to measure carbon emissions – often not open source, and more competitive than collaborative<br>No standardized way to report sustainability assessments in the US - lagging behind Europe where there is much broader alignment across different EU agencies<br>Uncertainty regarding continuity of government commitment to sustainability which makes it hard for companies to know what will stick.<br>Valley of death – hard to manufacture at scale, need more CMO type facilities, contract fermentation space<br>Lack of big infrastructure projects and investments that support the transition to new bio-based economy<br>Fuel markets are commoditized – prices are set and trades on global | Develop standardized metrics and measurement techniques<br>Explore developing a regulatory system for bio-based products and chemicals that is separate from their fossil fuel counterpart – which should be expedited base on risk assessment<br>Ensure/advocate for increased funding for regulatory agencies<br>Engage and collaborate with the private sector and support companies<br>Allocate funds for research aimed at enhancing our understanding of the circularity within the supply chain of various bio-based products.<br>Cost - biomanufacturing hubs could also help companies pilot these ingredients in a somewhat subsidized way initially<br>Set the market incentives to help create market certainty | "Circularity of the supply chain depends on what sort of circle you draw - for example, a company may only be sourcing what they consider "sustainable" sugar, but if you consider that that sugar would go into the food supply chain if it weren't for the fact that they're using it to make chemicals, you aren't getting the full picture because someone else might just be cutting down the rainforest to replace and plant that sugar that isn't going into the food supply."<br>"If your entire argument was saving the world from methane emissions off of cattle, and you have to feed 3 times more cattle in order to bring forth the alternative to milk, you aren't actually saving the world from methane. We need to be able to measure this."<br>"We need entities like BioMADE and other public entities trying to support a more holistic approach to |
|  | markets. Hard to compete with this.<br>Carbon-based products have benefited and continue to benefit from decades of large-scale investment and incentives.<br>Complex, patchwork regulatory landscape hinders progress | through standards<br>Pre-competitively reduce feedstock costs, making those more accessible and less energy and water intensive in order to produce - ultimately making the basic inputs to fermentation more accessible and more sustainable<br>Fund studies to measure crops best suited for CO2 capture and invest/support those efforts<br>Ensure policy consistency and longevity - needed for industry that is very capital intensive<br>Provide or design creative financial incentives, like loan guarantees for construction of facilities. | defining how we want to tackle measuring sustainability."<br>"One solution could be to separate industrial crops from food crops in the regulatory frameworks. But there is fundamental dilemma with that too: what happens when there is cross contamination across lands (when industrial crops contaminate food crops)? As our products move from greenhouses to real fields, will this become a bigger issue?"<br>"The fossil fuel incumbent has enjoyed a very substantial policy support system for a very long time."<br>"The reality is, we go through the same regulatory processes as our fossil counterparts - that's just the way it was historically set up. That was the easiest path to market. The reality is, we need a separate regulatory system for non-fossil building blocks and ultimately products. We are in the same queue as the products that the administration and society is looking to replace or make better. You can't have the impact on safety, security, social responsibility and sustainability if we're all moving in the same pipeline."<br>"We are not given any kind of increased standing compared to fossil fuel industry, and we should where there are fewer risks with bio-based products."<br>"Biotechnology just has such an enormous ability to impact and increase human health in particular around manufacturing sites, talking about being located around a petrochemical manufacturing site versus a |

|  |  |  | biotech or a bio manufacturing or fermentation site - the health impact and risks are just night and day.”<br>“I think a lot of people in this space are trying to do better with how they do the analyses for themselves, but challenges will persist until standardization is achieved”<br>“Making the products more sustainable isn't enough, we need to make the market more sustainable.”<br>“There needs to be instruments that are government led to provide economic incentives to make that transition.” |
| --- | --- | --- | --- |
| SOCIAL RESPONSABILITY |  |  |  |
| Perception | Barriers | Recommendations | Key Quotes |
| Social equity<br>Diversity and inclusion, including looking at economics and advancement<br>Environmental responsibility<br>Accessibility of the products which are more sustainable<br>Impact of new industrial spaces on frontline communities, including: land use interaction with local & indigenous communities<br>Transparency and traceability of supply chains<br>Bioethics<br>Job creation and job access<br>Avoiding monopoly<br>Cost parity vs benefit to society | Cost<br>Lack of standardized metrics to measure success<br>Lack complete, comprehensive, standardized definition<br>Lack of fast tracked or tailored regulations | Develop metrics and measurement techniques - social responsibility needs a global system of recognition and verification<br>Promoting engagement across different education streams<br>Gov program to improve cost accessibility of new green/sustainable products through tax incentives or other economical mechanisms<br>Promote/support sharing best practices<br>Making social responsibility part of the grant and loan application process, requiring grantees to describe how they are interacting with communities and how they thought about the potential impacts of their projects on the community<br>Increase and sustain investments in science and education so that all can benefit<br>Clarify confusion over Nagoya protocol and its enforcement (e.g., genetic material | “Maximizing benefits to mankind while minimizing the risks”<br>“Potential to create millions of high paying jobs”<br>“I really appreciate how it feels like a collaborative space of folks coming together rather than like a hyper competitive environment. That's been our experience like there's been really good knowledge, sharing community building through BioMADE in the past years.”<br>“It's important for the US to be a visible leader. There is an opportunity for the US to have significant influence, but we need to clean up our own backyard first.”<br>“Paramount to ensure we are including a diverse set of perspectives, is essential for creativity.”<br>“When you look at bio industrial manufacturing, it gives it a whole clean sheet. So, this gives us an opportunity to do this the right way and be welcoming to people of all race and |

|  |  | originating from Peru)<br>Help make opportunities, jobs, internships, etc. more visible and accessible<br>Support/Enforce Family policy/parental leave<br>Explore creative ways to increase access to new technologies (e.g., reassess patent longevity or include policies around ethical concerns in ROIs)<br>Involve companies of all sizes in discussions around policy making<br>Ensure the government is not sponsoring future monopolies | gender.”<br>“You have to ask yourself, which society? What do we mean by “social” exactly? And then you have to ask who's responsible? And then responsible for what? It gets messy very, very quickly in terms of what that actually means and what we owe to society writ large versus what we owe to just the advancement of knowledge and science.”<br>“There were issues with monopoly – production of grains owned by only 4 companies (90% of market), what can you do about this? And will the biomanufacturing ecosystem end up like this as well? Is the government funding something that will end in the hands of a few large corporations?”<br>“Every time you invent new tech, there is no guarantee it will be better. History shows that biotech improved significantly the prevention of hunger but it's still a big problem. So, it's still not equitable, even in a country like the USA with 38 million Americans living in poverty.”<br>“Social responsibility is not going to happen just by accident.”<br>“Many historical examples where technology is a form of oppression and has caused inequities, so that's still a public perception – we need to reckon on how to deal with that history” |
| --- | --- | --- | --- |
| EDUCATION & WORKFORCE |  |  |  |
| Perception | Barriers | Recommendations | Key Quotes |
| N/A | Talent retainment<br>Manufacturing job misperceptions<br>Siloes between academia, government, and private sector<br>No coordinated education | Develop workforce development incentives<br>Engage with traditional manufacturing pipeline<br>Increase K through 12 engagement and raise awareness about career path | “General workforce development incentives would be great. And this is something that obviously the Inflation Reduction Act and the EO on bio manufacturing have really focused in on. But we need |
|  | national standards | Develop/support personalized and dedicated mentorship programs in high school<br>Develop and share metrics and goals to guide education and training efforts<br>Invest in building biomanufacturing departments in community colleges<br>Increase investments in rural America for education programs<br>Increase job visibility and accessibility<br>Provide grants for students to support internships<br>Ensure that education program building and funding is bipartisan and consistent across administrations<br>Support the standardization of rational science education<br>Allocate funding to support initiatives aimed at gaining a deeper insight into workforce population dynamics, particularly in areas where there is a risk of an aging workforce<br>Engage with local and state economic development groups and state bio associations<br>Support the liberalization of our immigration policy<br>Support parental leave policy in the industry<br>Fund and develop more programs that allow for talent to gain experience in gov, academia and private sector and support talent transition in and out of different sectors (through coordinated advocacy, dedicated mentorship programs and targeted training programs) | to make sure that this is a career path that is attractive to people coming out of whether it's high school or even higher education.”<br>“The next set of true blue-collar jobs could be enabled by biology.”<br>“One doesn’t need to go to college to be to be operators of these facilities and get good high paying jobs.”<br>“The skills needed when defending your PhD are different to convincing a CEO about the value of a particular project – how do we teach people to be successful in such different environments and bridge the silos.” |
| **REGULATORY SPACE** |  |  |  |
| **Perception** | **Barriers** | **Recommendations** | **Key Quotes** |
| N/A | Patchwork of regulations (at the local, municipality, | Increase funding for further research to ensure that | “We do not go after alternative chassis, |
|  | state & federal levels)<br>Lack of human capital at regulatory agencies<br>Complexities around balancing safety to strengthen public trust and overregulation<br>No defined specific regulatory framework<br>Speed / backlog in application processes | regulations are grounded in sound scientific principles, rather than relying solely on linear extrapolation. Failing to do so may jeopardize the credibility of regulators.<br>Advocate and ensure that the regulatory system has the resources that it needs<br>Offer assistance to the private sector for the implementation of policies originating from the White House.<br>Develop a coordinated approach to engage government and private sector<br>Identify guardrails for BioMADE's potential engagement with Congress<br>Explore ways to allow for expanded access and harnessing of more natural biodiversity of organisms that can contribute to the biotechnology value chain.<br>Need to streamline/increase speed of processing time: much more rapid mechanism of product approvals<br>Explore a hierarchical approach to screening new materials as they progress from academia to prototyping to manufacturing.<br>Review how ingredients/components are categorized and how that influences the way they are regulated – sometimes not based in science (e.g., same product, different pathway))<br>Reevaluate how the current incentive structure and regulatory pathways may create inefficiencies by steering investments towards less useful ways to use a specific technology<br>Update a coordinated and rationalized framework to help innovation get to market | alternative microorganisms because of regulatory concerns.”<br>“Need to focus on the real threats, not the imagined ones.”<br>“We are coming in with a biologically produced replacement for something that is traditionally manufactured with petrochemicals, often with a manufacturing process that inherently involves multiple hazardous materials, whereas our bio-based process is a lot safer. It is a fermentation process, with carbohydrate, oxygen, mineral salt. But we still have to go through a regulatory framework that's based on something produced by something that's much more hazardous. It doesn't make sense.” |
| **COMPETITION** |  |  |  |
| **Perception** | **Barriers** | **Recommendations** | **Key Quotes** |

| N/A | Government historically not good at picking winners and losers<br>Patchwork regulatory landscape hinders competition<br>Cost of building manufacturing facilities<br>Cost of raw material cost & supply<br>– no competition on crop product buy-outs<br>Mammoth competitor lobbies<br>Valley of death for early commercialization | Government must be mindful of not creating a monopoly in their support of the industry, where only a few big companies end up dominating the market<br>Streamlining safety regulations will help more companies, especially smaller startups, compete<br>Create/Expand loan guarantee programs<br>Leverage national labs to assume an extended role as pre-competitive environments, modeling them after existing national renewable energy labs in Golden, CO. In such spaces, smaller groups could utilize the available facilities to assist with scale-up and conduct pilot studies, eliminating the need for them to invest their own funds in constructing facilities.<br>Develop industrial policy that will help bridge valley of death<br>Support increased security practices<br>Encourage people in the science, technology, and engineering community to participate in the policy development efforts | “Every year China graduates 1 million PhDs – how do you compete with that?”<br>“Bigger companies are better able to handle this complex regulatory patchwork because they have a regulatory team that can do that.”<br>“1 billion dollars funding – that’s a blip compared to the monolith of petrochemicals.”<br>“We’re too attentive to our IP concerns, and because we are so opposed to sharing tools and resources, our progress is really stymied. And it’s really disjointed, and much more expensive than it needs to be, in my opinion, than if we work together collectively on real opportunity spaces that can be truthfully precompetitive. Corporate entities can still get sufficient returns by demarcating where we can work up together to tackle some major challenges and hurdles that the industry is going to face.” |
| --- | --- | --- | --- |
| PUBLIC PERCEPTION |  |  |  |
| Perception | Barriers | Recommendations | Key Quotes |
| N/A | Public’s preconceived notions and fears<br>Historical context where new technology led to further inequities | Develop strategy to help industry preemptively communicate safety effectively with the public<br>Develop strategy to help industry communicate benefits to society/the community<br>Engage directly with communities to help build confidence in the technologies and products<br>Explore lessons learned from the GMO experience – start by promoting products that seem inherently non-threatening to get the public | “We could have the safest technology around. But if the community doesn’t get it, it’s not worth anything.”<br>“This will require a bit of a breakthrough in public thinking.”<br>“We need more people that are good at public communication and communicating science.”<br>“There’s always a risk and reward, and I think we do a disservice when we don’t present the risks associated with things.”<br>“People are very suspicious |
|  |  | exciting and educate public<br>on what biomanufacturing<br>can do<br>Focus on climate change<br>benefits in communication | of new technology” |

### Barriers & Recommendations

Participants identified barriers to the integration of the 4S Principles into the industry and outlined potential policy recommendations to address them (see table 3).

#### Safety

Many industry leaders highlighted the need for standardized metrics and evaluation methods to integrate safety principles effectively (e.g biocontainment measures or monitoring and recording of personal or environmental exposure incidents etc), emphasizing the importance of clear definitions and measurement techniques to assess safety compliance. One interview commented that “*because biotech is so fragmented, safety and security can mean different things to different people, and the significance of it can be different*”. Other leaders noted the complexity of risk assessments for emerging products, such as de novo proteins and systems. They urged increased investment in researching the safety and security implications of these technologies and the creation of updated risk models with supporting datasets.

#### Security - Biosecurity

A minority of interviewees were interested in biosecurity, defined as measures and practices employed to protect against the misuse of biological substances and technologies. They recommended the USG educate industry members about biosecurity, develop both educational and practical biosecurity tools, and encourage or mandate adherence to security best practices (e.g. screening DNA synthesis orders) through financial incentives or legislation. Some were concerned about the inefficiency of current gene or protein synthesis screening tools, as they may be defeated by AI-generated sequences. Some participants observed that understanding and assessing the interactions among different biological functions within a single cellular system is of crucial importance, one noted “*what do the systems do [biological] when put together? And could the outcome be weaponizable?”.* They emphasized however that there is a critical gap in expertise and tools for such assessment and prediction, which heightens biosecurity concerns, including predicting unintended consequences of such interactions and identifying potential dual-use research of concern (DURC).They urged the USG to invest in research for the safety and security of these technologies, develop updated models to predict biological functions for de novo biological products, and to consider strengthening the role of national Laboratories to address these challenges, one stated “*historically, the government had a responsibility to understand the safety and security implications of emerging technologies. I just don’t see that kind of like large scale investment going into the National Lab architecture*”.

#### Security - IP & Industrial Security

Most expressed concerns about IP security, calling for increased investments in genetically built-in protections which could thwart unauthorized use of a patented organism. Those technologies would address IP concerns with exports to foreign countries as well as bioterrorism concerns. Other industry leaders noted that as the economy moves towards more biomanufactured products, these facilities have the potential to become targets for IP theft; the USG should collaborate with companies to identify vulnerabilities to physical and cyberattacks and strengthen security across all production stages.

#### Security - National & Economic Security

A handful of industry leaders recommended the USG explore new export control policies, and clarify International Traffic in Arms (ITAR) and other foreign staff requirements for biotech and biomanufacturing; one stating “*we have multiple foreign nationals working on our project, and we’re doing our best to not share details beyond what they need to know. [the USG funder] never asked us to think about how we’re going to come up with checkpoints. I don’t really want more bureaucratic paperwork, but at the same time, it would be useful for them to start thinking about these potential concerns*”. Some advocated for policy research to understand the economic and geopolitical vulnerabilities associated with the bio-transition, including data infrastructures and supply chain visibility. They noted that such analysis is vital as it mirrors the strategic importance of managing critical resources like oil and gas. By mapping the sources of capabilities, infrastructure, and materials (e.g. feedstocks), as well as assessing related geopolitical risks, we can develop robust strategies to ensure sustainable, secure, resilient and equitable access to these key biological resources, mirroring strategies used in traditional energy sectors.

#### Sustainability

Bio-manufactured products are often associated with reductions in emissions, land-use, and water-use, but, most industry stakeholders emphasized that absent standardized reporting approaches and tools, generating comparable data is challenging, making it difficult for stakeholders to accurately assess and compare sustainability performance across different organizations or sectors. On participant stated “*I think a lot of people in this space are trying to do better with how they do their sustainability analyses, but it’s a challenge as it’s not standardized*”. Consequently, many suggested that the USG should play an active role in developing and standardizing sustainability metrics and benchmarks, particularly for carbon emission reporting and life cycle assessments, “*we need entities like BioMADE, and other public entities, to start supporting a more holistic reviewed approach to measuring sustainability*”. There was widespread agreement that standardized metrics are essential for consistent, transparent, and credible sustainability assessments, and would empower industry stakeholders to make more informed decisions, support effective policy development, and facilitate market access. Many stakeholders noted the dominance of carbon-based products due to historical investments and incentives, which makes it difficult for bio-based products to compete, as stated by one stakeholder “*the fossil fuel incumbent has enjoyed a very substantial policy support system for a very long time*”. Some recommended that the USG refocus investments on large-scale infrastructure projects to help the transition towards a bio-based economy. USG-sponsored manufacturing hubs could help companies pilot products through subsidization, accelerating product development and bridging the so-called “valley of death” where emerging pilot technologies often struggle to transition to large-scale production. Others thought market or other creative financial incentives (e.g., loan guarantees for construction of facilities; set standards) would promote market creation and drive further investments in the industry.

#### Social Responsibility

All industry stakeholders agreed that the bioeconomy has tremendous potential for equitable job creation across the country, including in disadvantaged or underrepresented communities, “*biomanufacturing has the potential to create millions of high-paying jobs”.* Many recommended the USG continue boosting domestic manufacturing jobs, albeit with a larger focus on bio-based manufacturing and equitable access to professional opportunities in this industry. Most highlighted that for biomanufacturing to deliver on its social responsibility promises, broader investments in science, technology, engineering, and mathematics (STEM) education are needed beyond the traditional graduate and postgraduate programs, including at the K1-12 levels, trade schools, and community colleges. Others suggested government should play a larger role to address diversity issues by co-funding student industry internships making these opportunities more visible and accessible across diverse populations. A handful of participants also called for more active and hands-on engagement with communities and local associations to better understand the impact of new technologies and new manufacturing hubs on their environment, lifestyle, and livelihoods.

#### Cross-Cutting Issues

Industry leaders emphasized that if the US is to be a global leader for the ethical standards and norms within bioindustrial manufacturing, it needs to be a global leader in bioindustrial manufacturing as a field. Thus,, it must address challenges impeding US competitiveness in this sector. They highlighted specific cross-cutting areas of interest fundamental to the success and advancement of the biomanufacturing industry, and for the successful integration of the 4S Principles.

#### Regulatory uncertainty

Most interviewees regarded the lack of a unified national regulatory framework for bioindustrial products as a significant barrier to establishing 4S standards across the industry – one stated “s*afety regulations are completely inconsistent, so everybody gets stalled out*” and another “*there’s really terrible safety regulation uniformity on engineered organisms in the US*”. State-by-state approaches, for example in the regulations of genetically engineered crops, and other regulatory disparities among various agencies further complicate matters, affecting public clarity, trust, and the ability of smaller companies and startups to innovate, compete, and expand nationally. Many suggested streamlining the regulatory processes would help bioindustries compete fairly against fossil fuel-based technologies, which have historically enjoyed substantial policy and financial support, “*we go through the same regulatory processes as our fossil counterparts. The reality is, we need a separate regulatory system for non-fossil building blocks and ultimately products. We are in the same queue as the products that the [Biden] Administration, and society as a whole, is looking to replace or make more sustainable. We can’t have the impact on safety, security, social responsibility, and sustainability if we’re all moving in the same pipeline*”. Some called for a separate, expedited regulatory pathway, based on robust risk assessments, so that sustainable products could more easily enter markets. some technologies do not have a clear pathway at all, including organisms for remediation, which contribute to delays and bottlenecks in these technologies becoming available. Increased resources and expertise within US regulatory agencies were recommended to expedite reforms to facilitate competitiveness.

#### Education and workforce development

A robust biomanufacturing workforce is needed for the success of the industry; participants cited skilled staff shortages, talent retention issues, and workforce preferences as factors exacerbating a critical workforce deficit. Others noted that silos between academia, government, and the private sector hinder knowledge exchange, creating a skills gap, and that the absence of standardized national guidelines leads to inconsistency in educational programs. There was widespread agreement that specialized scientific knowledge from advanced education programs is no longer a necessity in the industry, and that bachelor’s and associate degrees, as well as certificates, could adequately train most workers, and that these training and hiring trends must accelerate. One participant astutely stated “*the next set of true blue-collar jobs could be enabled by biology*”. Interviewees suggested government actions to foster a skilled and sustainable biomanufacturing workforce, including implementing workforce development incentives, increasing government funding in K1-12 STEM education and revising curricula to align more closely with industry demand. To expand access to training, many called for funding community college biomanufacturing programs and provide grants for student internships that help create equal opportunities for practical experience and potential future employment in the biomanufacturing sector. One stakeholder said “*general workforce development incentives would be great. This is something that obviously the Inflation Reduction Act and the executive order on biomanufacturing have really focused on. But we need to make sure that this is a career path that is attractive to people coming out of, whether it’s high school, or even higher education, and showing the possible career development*”. Other leaders noted that targeted immigration reforms and family-friendly policies, like parental leave, attract skilled workers, enrich the talent pool, and demonstrate commitment to a more inclusive workforce.

#### Public Perception

Several identified public perceptions of bio manufactured products as a key risk factor for the overall success and flourishing of the industry, with one participant stating “*we could have the safest technology around. But if the community doesn’t get it, it’s not worth anything*”. They recommended the USG take strategic actions to avoid public misperceptions of the technology through lessons learned from genetically modified organisms (GMO). Communications could highlight the societal benefits of bio-based technologies, especially regarding climate change and the economy. Some recommended a communication strategy focused on products with widely accepted benefits, that replace products that rely on animal or petrochemical inputs. Many noted that engaging directly with communities is essential to build public trust and support in biotechnologies and products.

#### Competition

Many stakeholders highlighted the importance of making the industry more competitive as a critical step towards integrating the 4S Principles but cited several barriers hampering these efforts. The influence of powerful competitor lobbies, including petrochemicals, was named by most participants as a critical barrier to the industry’s advancement, with one stakeholder stating “*1 billion dollars funding… that’s a blip compared to the monolith of petrochemicals*”. Others cited the fragmented regulatory landscape as major hurdle for innovation and competitiveness, while some highlighted the high cost of building biomanufacturing facilities, coupled with limited competition in raw material supply, as major challenges for smaller companies and start-ups. Some interviewees suggested that the USG expand loan guarantee programs and leverage National Labs as pre-competitive hubs, to facilitate scaling and pilot studies without extensive private investment. The majority also agreed that it was critical for policymakers to involve the science, technology, and engineering communities in policy development to better support the biomanufacturing industry’s growth competitiveness.

## Discussion

In response to the potential global technology race, global powers like China, the UK, and the EU are rapidly expanding their biomanufacturing capacity and workforce. China, in particular, is narrowing the gap with the U.S. through aggressive R&D investments, targeted government directives under their “Made in China 2025” plan, and a strong focus on STEM talent attraction and retention.(*4, 5*) President Biden’s Executive Order 14081, issued on September 12, 2022, initiated the “National Biotechnology and Biomanufacturing Initiative”, which outlines a whole-of-government approach to propel biotechnology and biomanufacturing towards innovative solutions in various critical areas, such as health, climate change, energy, food security, agriculture, supply chain resilience, and national and economic security. Meeting the EO’s ambitious targets would not only establish a solid basis of U.S leadership and competitiveness in biobased manufacturing but would also position it to lead the shaping of international ethical standards and norms.(*6*)

The interviews conducted in this study yielded valuable recommendations for the integration of the 4S principles into the industry at large. However, a key takeaway emphasizes that for the U.S to truly become a standards setter rather than a standard taker, it must first secure its position as a competitive leader. Industry stakeholders have emphasized several overarching barriers that must be addressed to facilitate the growth of the U.S. bioindustrial economy. These barriers include a fragmented and redundant regulatory landscape, deeply entrenched policies and incentives that still benefit the petrochemical incumbent, as well as significant workforce shortages. To successfully transition and establish itself as a global leader in the bioeconomy, the U.S requires a comprehensive, government-wide strategy aimed at the radical transformation and restructuring of its industrial complex.

In 1986, the EPA, FDA, and USDA jointly published the U.S. Coordinated Framework for Biotechnology Products, which was updated in 2017.(*7*) This framework offers guidance and regulations for biotechnology products but does not address the increasing complexity of emerging bioproducts that often do not operate within a single agency’s jurisdiction. This lack of clarity often results in companies having to file regulatory documents with multiple agencies, using different data and formats, leading to inefficiencies for both regulators and product developers. Stakeholders interviewed in this study suggest streamlining regulatory processes to enhance competitiveness against the fossil fuel-based incumbent, which historically received substantial policy and financial support.(*8*) Some proposed an expedited regulatory pathway, based on robust risk assessments, to facilitate market entry for bio-based products which are inherently more sustainable. Others proposed a science-based approach to streamlining safety regulations, reducing redundancy in safety testing like animal testing of established chemicals, and suggested exploring chemical equivalent reach-across programs to boost competitiveness and expedite market entry for biomanufacturing products. The President’s Council of Advisors on Science and Technology provided complementary recommendations in their own report to the President. Specifically suggesting that regulatory agencies should draft updated and streamlined model pathways for review of emergent bioproducts, as well as establishing and training an inter-agency rapid response team to vet new crosscutting bioproducts.(*5*) There may also be an opportunity for regulatory authorities to harness the potential of AI and advanced data analytics to streamline regulatory processes, bolstering efficiency and reducing bottlenecks. As other nations expedite their regulatory review and approval procedures, it is now more critical than ever for the U.S. to address its regulatory bottlenecks to maintain its competitiveness and leadership role.

Industry leaders interviewed in this study all agreed the industrial biorevolution had the potential to create millions of high paying jobs across the country, including in economically and socially marginalized communities. A strong, well-trained and diverse workforce is also necessary to manifest the bold promises outlined in the presidential EO and will require federal and state governments to offer incentives that facilitate collaboration between various educational institutions, including community colleges, Historically Black Colleges and Universities, Tribal Colleges and Universities, traditional 4-year universities, as well as K-12 schools. Such programs would help develop robust curricula and certification programs in biomanufacturing science and would ultimately help foster diversity by equipping individuals with the in-demand skills needed for immediate employment in the biomanufacturing industry.

In rural areas and regions rich in sustainable biomass resources, fostering the development of biotechnology capabilities through investments in training programs and local infrastructure not only has the potential to create new employment opportunities for these communities, but also to fuel the expansion of the U.S bioeconomy.(*9*)

It is clear that biomanufacturing will have significant impacts on carbon emissions, economic opportunities, as well as national security. Promoting and integrating the ethical principle in the biotech industry has long been an area of U.S policy development. However, this study marks a seminal shift by incorporating the viewpoints of biotechnology practitioners into these policy discussions and by broadening their scope through the inclusion of nontraditional bioethics issues like sustainability. While various models have been examined to integrate social and ethical expertise into technology development, the valuable insights from this study are poised to help reshape future US government policymaking. More importantly, this study underscores that for the U.S. to become the global leader in establishing standards and norms, it must first address the obstacles hindering the full realization and growth of its Bioindustrial sector.

## Limitations

While this study employed a robust methodology, certain limitations must be acknowledged. The inclusion of a diverse range of BioMADE members was a strength, but not all members could be interviewed. Despite conducting interviews with strict not-for-attribution policies and confidentiality measures, concerns around competitiveness may have influenced overall participation. Finally, it is important to note that the statements and recommendations provided by interviewees may not necessarily align with their organization’s official stance.

## Conclusion

With rapid developments in biotechnology and biomanufacturing, there is a pressing need to create fresh frameworks for incorporating BioMADE’s 4S Principles into the development of these new technologies. Moreover, a narrow window exists for the US to help promote impactful new norms as the field continues to expand globally, ultimately enhancing public acceptance and the success of industrial biotechnology. The insights, concerns, and recommendations detailed in this study are poised to inform future USG policymaking and norm-setting, shedding light on the crucial perspectives of industry practitioners.

## Acknowledgments

We would like to thank all the study participants for their time and valuable insights, as well as the BioMADE leaders, especially Beth Vitalis, for their ongoing support throughout the project.

## Author contributions

Methodology: Aurelia Attal-Juncqua and Gigi Gronvall

Investigation: Aurelia Attal-Juncqua, John Getz, Ryan Morhard and Gigi Gronvall

Funding acquisition: Gigi Gronvall and Ryan Morhard

Supervision: Gigi Gronvall

Writing – original draft: Aurelia Attal-Juncqua

Writing – review & editing: Aurelia Attal-Juncqua, John Getz, Ryan Morhard and Gigi Gronvall

## Data and materials availability

All data are available in the main text or the supplementary materials.

